# Active Remote Focus Stabilization in Oblique Plane Microscopy

**DOI:** 10.1101/2024.11.29.626121

**Authors:** Trung Duc Nguyen, Amir Rahmani, Aleks Ponjavic, Alfred Millett-Sikking, Reto Fiolka

## Abstract

Light-sheet fluorescence microscopy (LSFM) has demonstrated great potential in the life sciences owing to its efficient volumetric imaging capabilities. For long term imaging, the light-sheet typically needs to be stabilized to the detection focal plane for the best imaging results. Current light-sheet stabilization methods rely on fluorescence emission from the sample, which may interrupt the scientific imaging and add to sample photobleaching. Here we show that for oblique plane microscopes (OPM), a subset of LSFM where a single primary objective is used for illumination and detection, light-sheet stabilization can be achieved without expending sample fluorescence. Our method achieves ~43nm axial precision and maintains the light-sheet well within the depth of focus of the detection system for hour**-**long acquisition runs in a lab environment that would otherwise detune the system. We demonstrate subcellular imaging of the actin skeleton in melanoma cancer cells with a stabilized OPM.

## 1. Introduction

Light-sheet fluorescence microscopy (LSFM) has emerged as a fast and gentle method to acquire volumetric image data [1]. As such, LSFM has found various applications ranging from single-molecule imaging to longitudinal imaging of model organisms. In a typical LSFM implementation, the illumination and detection are implemented through separate objective lenses, oriented at 90 degrees to each other [1]. The system is then carefully aligned such that the light-sheet plane overlaps with the focal plane of the detection system. As such, if the detection objective experiences focal drift (for example through thermal expansion [2]), the relative alignment of the light-sheet and the detection is altered, and if left unchecked, the light-sheet can drift outside the depth of focus of the detection system. As a result, the imaging performance of the LSFM is drastically reduced, as only out-of-focus blur is collected. Several “auto-focusing” algorithms have been developed for LSFM [3-5], which typically acquire a series of fluorescence images to estimate the relative displacement. This requires the expenditure of fluorescence for such measurements, and an interruption of the actual imaging operations. This is different to autofocusing systems in conventional, epi-fluorescence microscopes, which typically measure axial drift from a laser beam that is back-reflected from the coverslip [6-8]

Oblique plane microscopy (OPM) is a variant of LSFM which uses a single objective lens to launch the light-sheet at an oblique angle and detects the fluorescence with the same lens [9, 10]. As such, if the primary objective lens exhibits drift, the light-sheet and detection drift at the same rate, and their relative alignment stays intact. The alignment is only broken for higher order aberrations (i.e. imaging in inhomogeneous media), but can be restored via adaptive optics in an OPM [11].

In a typical OPM, the tilted light-sheet plane is mapped through a remote focusing system [12] onto a camera, which consists of a secondary and tertiary objective. Any drift of the secondary and tertiary objective lenses will shift the detection focal plane, but not the light-sheet. As such, the remote focusing system can cause a similar image degradation as in a conventional LSFM that experiences light-sheet drift.

Here we show that in an OPM, the remote focusing unit can be stabilized with laser light, which can be spectrally distinct from the fluorescence detection. The system does not need fluorescence light for light-sheet stabilization and runs continuously and independently to fluorescence imaging. This contrasts with light-sheet stabilization in conventional LSFM architectures, which must be interlaced with fluorescence imaging operations. We characterize the precision of the system and demonstrate hour long high resolution OPM imaging of subcellular dynamics.

## 2. Methods

### 2.1 Optical setup

In Figure 1A, our implementation of OPM is shown. For illumination, laser light from a previously published light-sheet module [13] is coupled over a dichroic mirror (Di03-R405/488/561/635-t1-25×36, Semrock) into the optical train of the OPM. After passing through two tube lenses (TL 1: ITL-200, TL2: custom, 150mm EFL) and two galvo mirrors (Pangolin Saturn 9B with a 10mm y-mirror mounted 90 degrees rotated) for image space scanning [14], an oblique sheet is launched in sample space through the primary objective (O1, Nikon 40X, NA 1.25 silicone oil).

**Fig. 1.**
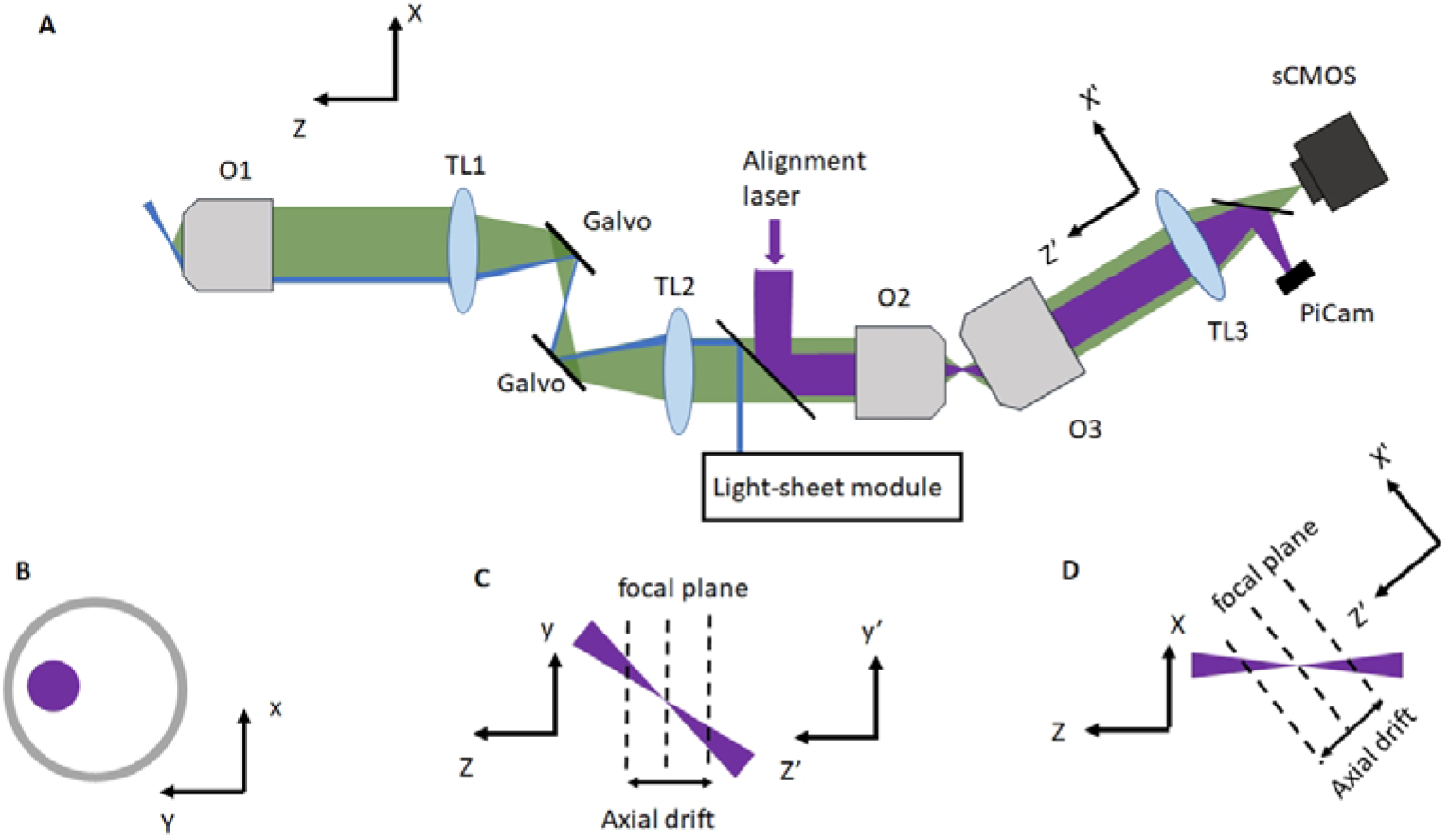
Schematic layout of an oblique plane microscope with remote focus stabilization. **A** Schematic layout of the OPM. Green shows the fluorescence path, blue shows the light-sheet path and magenta shows the alignment laser path. O1: primary objective, O2: secondary objective, O3: tertiary objective. TL1-TL3: first, second and third tube lens. **B** Position of the alignment laser beam (blue) in the pupil of the secondary objective O2. **C** Laser focus in the remote imaging space between O2 and O3. **D** Laser focus shown in an orthogonal plane to **C**. Dotted lines correspond to the focal plane of O3.

Fluorescence light is detected through the same objective, whose pupil is conjugate to the secondary objective (O2, Nikon 40X NA 0.95). The secondary objective maps fluorescence emitters along the light-sheet into the remote space while minimizing spherical aberrations. A tertiary imaging system is used to map the fluorescence on a scientific CMOS camera (Orca Flash 4, Hamamatsu). The tertiary imaging system consists of a glass-tipped objective (AMS-AGY v2 54-18-9, Applied Scientific instrumentation) and a tube lens (f=300mm achromatic doublet, Thorlabs).

For the remote focus stabilization, we injected a collimated laser beam over the backside of the dichroic mirror into the optical train of the OPM (Figure 1A). For the experiments shown here, we used a fiber coupled 488nm laser (Edmund Optics, 10mW Pigtailed Laser Diode, part nr: #23-761), which was collimated with a f=100mm lens. The reflective side of the dichroic faces towards the primary objective (i.e. the light-sheet illumination laser bounces off the coated front surface of the dichroic). In contrast, the alignment laser first travels through the dichroic mirror substrate. As such, there is a secondary reflection from the substrate itself. We reasoned that if we use a laser line for which the dichroic was optimized for reflection, the secondary reflection would be minor. Indeed, we observed a notable double reflection when using another wavelength (i.e. laser diode at 785nm), whereas the 488nm line showed a single dominant reflection.

The alignment laser beam passes through the secondary and tertiary objective and is picked up by a dichroic mirror (Semrock Di02-R488-25×36) after the tertiary tube lens. The laser beam is then focused on a camera (OV9281-120, labeled PiCam in Figure 1A) used for the focus stabilization feedback. While it is advisable to put filters and mirrors in the infinity space, we found that with a tube lens of low optical power, placing the dichroic in the “image space” causes negligible aberrations. Adopters of the technology may consider placing the dichroic in the infinity space of the tertiary imaging system when using a tube lens with shorter focal length.

Our system measures the relative misalignment of the remote focus system by using a laser beam that is tilted to the optical axes of the secondary and tertiary objective (labeled Z and Z’ respectively). As such, an axial misalignment not only causes the laser beam to defocus, but also to be translated on the alignment camera. To this end, we inject the alignment beam at an off-center position into the secondary pupil (Figure 1B). This causes the beam to tilt in the remote focus space, as shown in Figure 1C. As such, if either the secondary or tertiary objective is shifted axially, i.e. its focal plane moves, the beam is defocused, but also translated laterally along the Y’ axis on the alignment camera.

In the other dimension, a tilt of the laser beam occurs naturally, as the tertiary imaging system is angled to the optical axis of the secondary objective (Figure 1D). In case of focal drift of the tertiary objective, the laser beam gets translated in the X’ direction, whereas a drift from the secondary objective will not show in the X’ direction (see also Figure 1D for an illustration).

The off-centering of the alignment laser in the O2 pupil increases the sensitivity of the measurement (a given focal shift causes a larger lateral shift on the camera), but it also ensures that axial shifts of both O2 and O3 become measurable. To place the laser beam off-center in the pupil of O2, its numerical aperture has to be reduced (i.e. the laser beam underfills the pupil). This in turn increases the depth of focus of the laser beam and hence makes fitting of the laser spot easier over a larger defocus range. Importantly, it does not lower the localization precision for the scenario of “unlimited photons”. While the spot gets larger in size when reducing its NA, more photons (if the laser intensity of the laser beam is increased accordingly) contribute to the measurement, which restores the measurement precision.

### 2.2 Feedback Control

The feedback control for the remote focus stabilization was implemented on a Rasperry Pi 4B architecture building on the previous PiFocus [6] Python code. The Raspberry Pi continuously acquires a stream of images at 100fps from the PiCam, processes them in real time to compute an error signal of the laser spot, and uses this signal to control the axial position of the tertiary objective via a piezo stack actuator (PC4GR, Thorlabs). The Piezo stack was fitted with a custom mount into a manual 3D stage (Newport 9063-XYZ), where it rests against a differential micrometer.

We implemented a proportional-integral-derivative (PID) controller for fine-tuning the laser spot position and ensuring stabilization. The proportional (P) and integral (I) gains were experimentally optimized to minimize oscillations and suppress displacements of the laser spot. In this implementation, the derivative term was initially set to zero to simplify tuning and prevent overshoot, although it remains an option for future enhancements. The error signal is computed as the Euclidean distance between the current position of the laser spot and its initial calibrated position (setpoint). The position of the laser spot is determined by Gaussian fitting of the 1D projections of the laser intensity distribution (point spread function, PSF) on the x’ and y’ axes. By summing projections along the x’ and y’ axes and fitting 1D Gaussian profiles to the data, the system achieves reliable sub-pixel localization of the laser spot even under varying intensity conditions.

For each frame, the PID algorithm operates as follows:

1. Error Calculation:

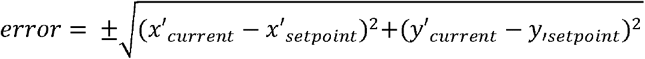 The sign of the error depends on the relative position of the laser spot on the camera sensor, ensuring directional correction.
2. Control Signal Computation

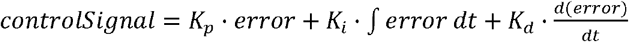
  ∘ The proportional term (*K*_p_) corrects the displacement magnitude.
  ∘ The integral term (*K*_i_) accumulates the error over time, reducing steady-state offset.
  ∘ The derivative term (*K*_d_) predicts future errors based on the rate of change, though it was not used in the current implementation to prevent noise amplification.
3. Actuator Update:

The control signal is translated into a voltage adjustment for the piezo stack, ensuring precise axial positioning of the tertiary objective. The corresponding conversion was found by measuring a calibration curve (see also Figure 2B).

**Fig. 2.**
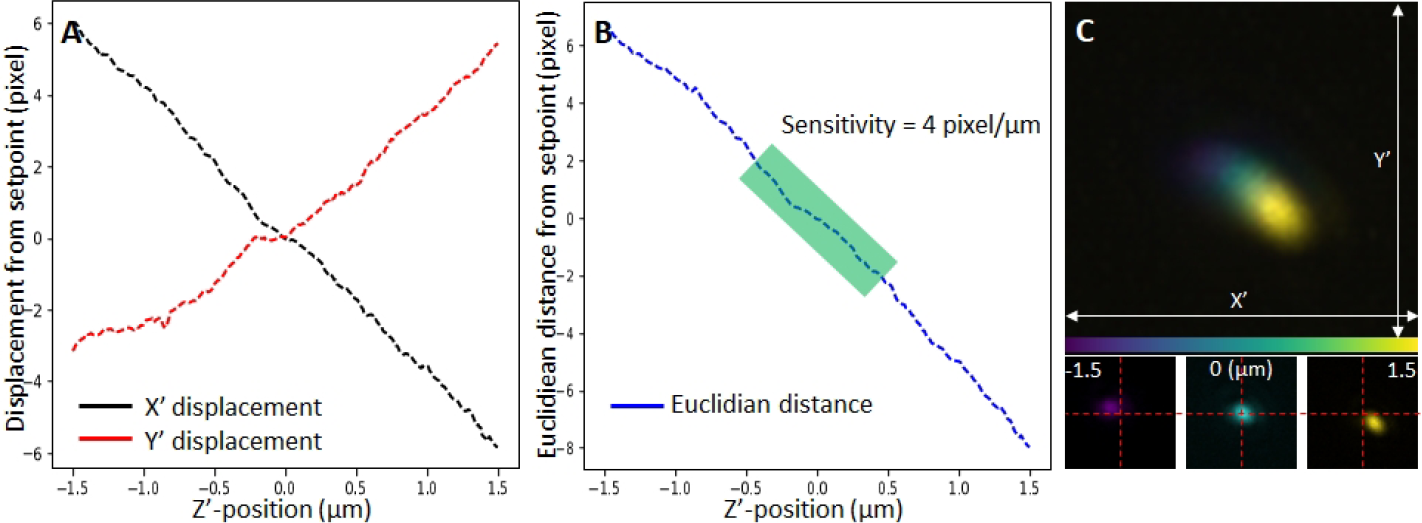
Axial drift estimation using spatial displacement. **A** X’ and Y’ displacement of the laser PSF extracted from tertiary objective Z’-scan. **B** Calibration curve of focal drift estimation using Euclidean Distance. **C** Z’-position color-coded projection image of 300 laser spots captured from a tertiary objective scan, insets show the PSFs at -1.5-micron, 0 micron and +1.5 microns. The laser PSF location moves diagonally over the image sensor through the Z’-scan.

We wrote a GUI which allows a user to calibrate the system at the start of an experiment (see also the note in the Appendix). In this calibration step, the system captures the initial laser spot position, defining it as the setpoint. Subsequent deviations from this position determine the error signal. The GUI also allows users to adjust critical parameters such as PID gains and verify system alignment. During operation, the system logs the error, control signal, and actuator position for each frame. This data is stored in memory and can be saved as a CSV file for post-experiment analysis. Real-time feedback allows users to monitor the control signal and ensure proper alignment throughout the experiment.

## 3. Results

### 3.1 Precision and long-term stability

To measure the accuracy of our stabilization method, we acquired a rapid timelapse sequence, with a duration of 100 seconds to minimize drift. The standard deviation of the localization measurement of the laser spot was then converted into nanometers using a calibration curve (where the piezo stack was stepped in known increments), as shown in Figure 2A-C and Figure 3A. We measured a standard deviation of ~57 nm using this approach. If converted back to the sample space (demagnification of 1.33), this corresponds to an axial precision of 43 nm, which is an order of magnitude lower than the depth of focus of the detection system.

**Fig. 3.**
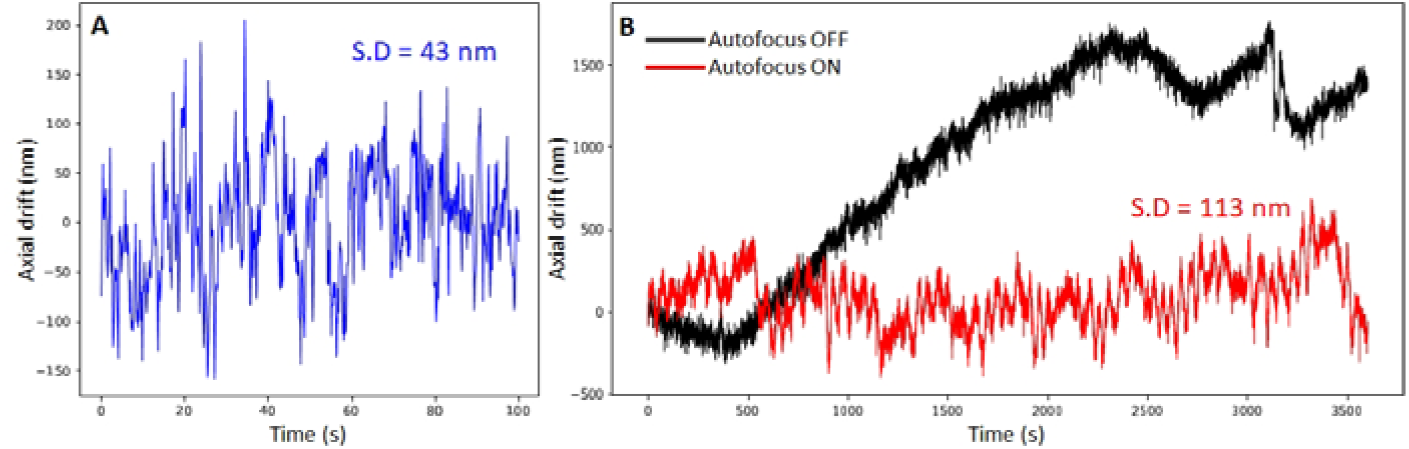
**A** Time-lapse acquisition at focal plane for estimating axial precision. **B** Axial drift with stabilization ON (red) and OFF (black) monitored for 1 hour. S.D. Standard deviation.

We then compared the long-term stability of the remote focusing system with and without focus stabilization. Without feedback correction, the system drifted ~ 2 microns over the course of an hour (Figure 3B). This is not unexpected, as our laboratory experiences temperature oscillations with an amplitude of ~1 degrees Celsius (see also Appendix for temperature measurements). With the focus stabilization on, the standard deviation over an hour was 150 nm measured in the remote space, corresponding to 113 nm mapped into sample space.

### 3.2 Imaging of fluorescent nanospheres and melanoma cancer cells

To test the effectiveness of our remote focus stabilization system during OPM imaging operations, we first volumetrically imaged 100nm diameter fluorescent nanospheres over one hour. In Figure 3 A-B, the first and the last timepoint of volumetric imaging are shown, with the stabilization system turned off. At the beginning of the run, we manually adjusted the light-sheet in respect to the detection for best imaging performance, yielding quite compact point spread functions with little out-of-focus blur (Figure 4A). As evidenced in Figure 4B, imaging performance deteriorated after one hour, as the light-sheet no longer overlapped with the detection focal plane, and mainly out-of-focus blur is excited.

**Fig. 4.**
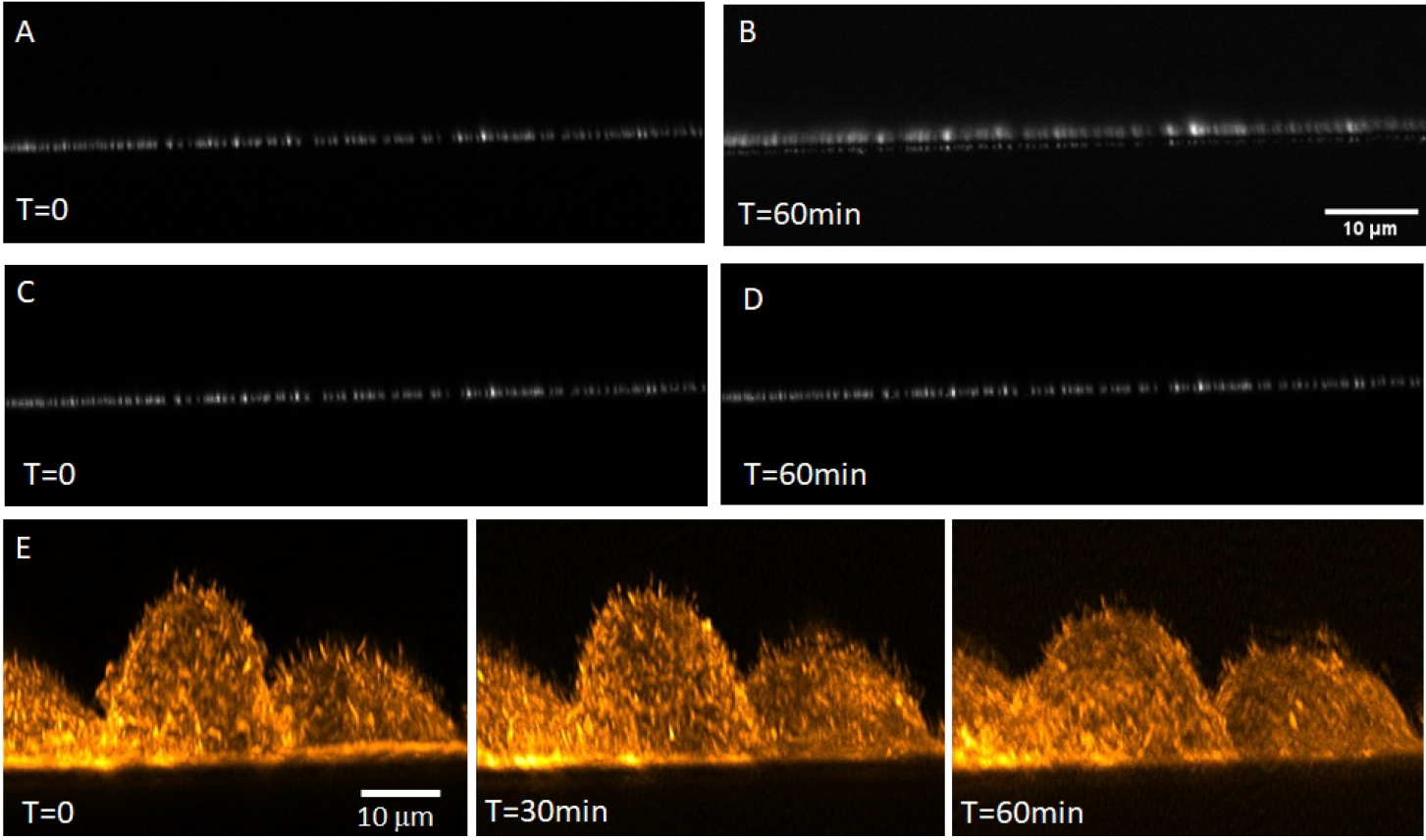
Time lapse volumetric imaging using OPM with and without remote focus stabilization. **A-B** 100nm fluorescent nanospheres, as imaged with OPM without remote focus stabilization at the beginning and end of an one hour time lapse series. Cross sectional maximum intensity projections are shown. **C-D** OPM imaging of the same volume as shown in **A-B**, but with remote focus stabilization active. **E** Three timepoints of a timelapse of A375 cancer cells labeled with tractin-mRuby, using OPM with remote focus stabilization active. Cross sectional maximum intensity projections are shown.

In Figure 4C-D, a second time-lapse imaging experiment is shown, with the stabilization system turned on. The system remained well aligned during the run, as evidenced by the last timepoint shown in Figure 4D.

Encouraged by these results, we performed volumetric time-lapse imaging of A375 cancer cells labeled with tractin-mRuby over one hour (2-minute time interval, for a total of 30 volumetric acquisitions) with the stabilization system turned on. Cross-sectional maximum intensity projections at three time points are shown in Figure 4E. The data was computationally deconvolved, and the contrast was manually adjusted for each image to compensate for the effect of photobleaching. Fine filopodia are visible on the three cells and remain well resolved during the entire course of the time-lapse acquisition.

## 4. Discussion

We demonstrate a simple remote focus scheme that can be implemented in OPM systems with minor modifications. Our scheme greatly increases the long-term performance of an OPM, and compared to other light-sheet modalities, stabilization does not interrupt the imaging operations, nor does it dispense fluorescence.

Improvements to our stabilization method could involve adding PSF engineering to the laser spot. In particular, making the depth sensing independent on the position of the spot on the camera could counter sources of long-term drift. As an example, O3 is mounted on a 3D manual translation stage. If any of its lateral axes (along X’ and Y’) drift, this would cause a translation of the laser spot on the camera, which could be mis-interpreted as a focal shift. We think that such other sources of drift may cause the larger standard deviation on our hour-long measurements. For the imaging experiments presented herein, long term stability on the order of 100nm are acceptable. However, if more complex light-sheets generated by optical lattices[15], Airy beams [16] or Field Synthesis [17] are used, potentially a higher precision could be desirable, especially for deconvolution. We estimate that by using a higher grade camera with a larger dynamic range than the PiCam used here could improve the localization precision by 2-5 fold.

We did not fix any drift of the primary objective. For volumetric imaging some small axial drift can be tolerated; It simply shifts the volume that is being imaged, without noticeably affecting the imaging point spread function. For instance, the fluorescent nanospheres and cells shifted a few microns downwards in Figure 4 over the course of an hour. As long as the sample stays within the beam waist of the light-sheet, this drift does not noticeably deteriorate imaging performance. Nevertheless, a secondary focus stabilization [6] for the primary objective could be a valuable addition to an OPM.

Overall, we hope that this resource will find widespread applications in OPM and will enable higher quality data acquisitions over extended time periods.

## 5.1 Acknowledgment

We would like to thank Dr. Vasanth Siruvallur Murali for preparing the A375 cancer cells. R.F. is thankful for support from the National Institute of Biomedical Imaging and Bioengineering (grant R01EB035538) and the National Institute of General Medical Sciences (grant R35GM133522).

## 5.2 Code availability

The python code for the remote focus stabilization can be accessed here: https://github.com/AdvancedImagingUTSW/OPM-Autofocus

## Appendix 1: Graphical User Interface (GUI) for Autofocus Control System

The GUI designed for the autofocus system serves as an interactive platform for setting parameters, monitoring performance, and visualizing the point spread function (PSF) in real time. It is built using the Tkinter library in Python and is optimized for user-friendly operation with a touchscreen. This appendix provides a detailed description of the GUI layout and the functions of each component.

### 1. Main Layout and Frames

The GUI is organized into multiple sections for clarity and functional grouping. The groups are dynamically sized to accommodate user inputs and visual feedback.

**Fig. S1.**
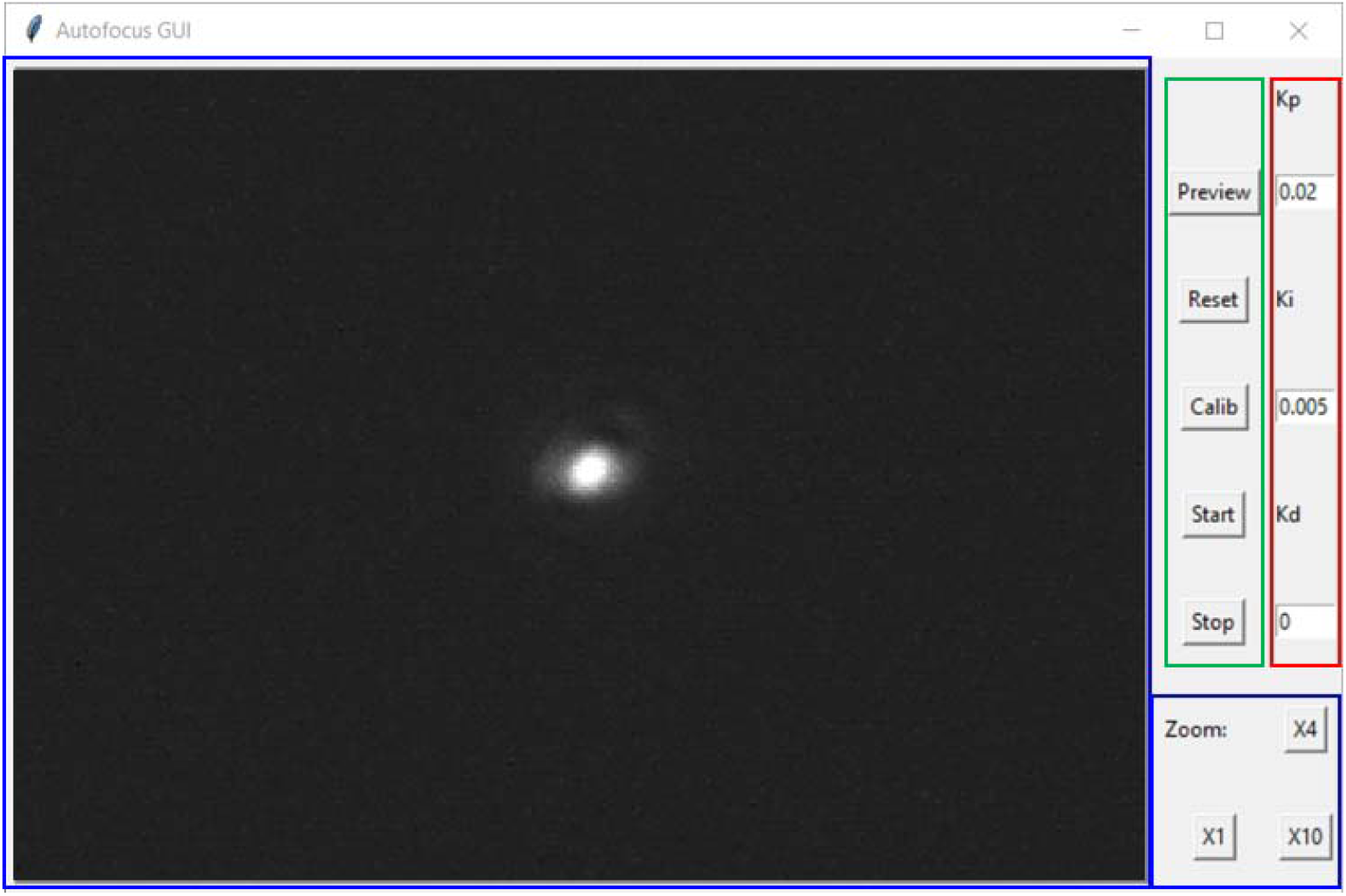
Graphical User Interface (GUI) for Autofocus Control System

#### a) Image Preview Group (blue box in the Fig. S1)

This group provides a live feed of the camera view, enabling visualization of the PSF in real-time.

Image Canvas:

A canvas widget where the live feed from the PiCam is displayed. This feed includes the real-time PSF overlays with the calculated center and setpoint for quick verification.

Zoom Controls:

1x 4x and 10x buttons to zoom in on a region of interest (ROI) for detailed observation of the PSF

#### b) Control Parameters Group (red box in the Fig. S1)

This group is dedicated to parameter input for the autofocus system. It allows users to fine-tune the PID control loop and other system settings.

Proportional Gain (P):

A text box where users can input the proportional gain (Kp) for the PID controller. This controls the immediate response to error.

Integral Gain (I):

A text box for entering the integral gain (Ki), which addresses cumulative error over time.

Derivative Gain (D):

A text box for setting the derivative gain (Kd). Although Kd is initially set to zero, this option allows flexibility for future adjustments

#### c) System Calibration Frame (green box in the Fig. S1)

This group allows users to calibrate the system before starting an experiment. Calib Button:

A button to initialize the laser spot’s reference position (setpoint). This ensures accurate error calculations throughout the session.

Reset Button:

A button to reset the calibration and clear the setpoint, allowing for a new reference point to be defined.

Start/Stop Buttons:

Buttons to start/stop the PID autofocus loop.

Preview Button:

A button to toggle the preview function.

### 2. Interaction Workflow

**Setup**:

The user inputs PID gains in the **Control Parameters Group**.

The **Calibration Button** in the **System Calibration Group** is pressed to set the initial position of the laser spot.

**Operation**:

The real-time feed in the **Image Preview Group** provides visual feedback of the PSF.

**Adjustments**:

Parameters can be updated: click **Stop Button** -> adjust PID gains in the **Control Parameters Group** -> click **Start Button**.

The user can recalibrate the system if necessary: click **Stop Button** -> click **Reset Button** and repeat the **Setup** process to set a reference position (setpoint).

## Appendix 2: Temperature measurements

We attached a temperature probe to the tertiary objective and measured the temperature over ~four days, as shown in Figure S2.

**Fig. S2.**
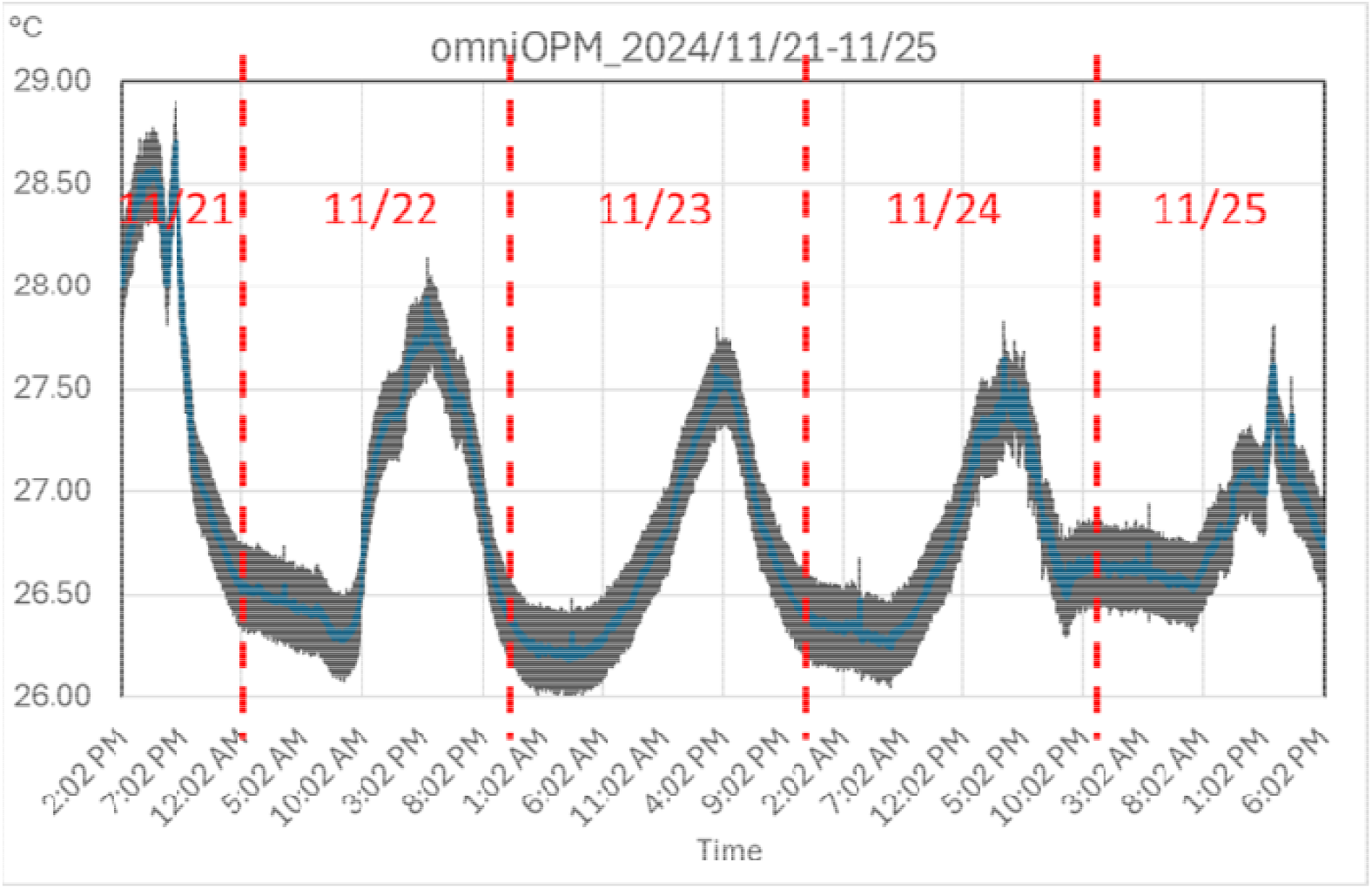
Temperature measurement on the tertiary objective.

The temperature oscillates during the daytime by 1.5-2 degrees Celsius peak to peak. Most experiments were done either in the morning or late afternoon, which saw strong temperature gradients. The only stable temperature period occurred after midnight, but no imaging experiments were performed at that time.

